# Tissue-specific knockout in *Drosophila* neuromuscular system reveals ESCRT’s role in formation of synapse-derived extracellular vesicles

**DOI:** 10.1101/2023.09.25.559303

**Authors:** Xinchen Chen, Sarah Perry, Bei Wang, Shuran Wang, Jiayi Hu, Elizabeth Loxterkamp, Dion Dickman, Chun Han

**Affiliations:** Weill Institute for Cell and Molecular Biology and Department of Molecular Biology and Genetics, Cornell University, Ithaca, NY 14853, USA; Department of Neurobiology, University of Southern California, Los Angeles, CA 90089, USA

**Keywords:** CRISPR/Cas9, tissue specific, CRISPR-TRiM, neuromuscular junction, SNARE, ESCRT, extracellular vesicles, *Drosophila*

## Abstract

Tissue-specific gene knockout by CRISPR/Cas9 is a powerful approach for characterizing gene functions in animal development. However, this approach has been successfully applied in only a small number of *Drosophila* tissues. The *Drosophila* motor nervous system is an excellent model system for studying the biology of neuromuscular junction (NMJ). To expand tissue-specific CRISPR to the *Drosophila* motor system, here we present a CRISPR-mediated tissue-restricted mutagenesis (CRISPR-TRiM) toolkit for knocking out genes in motoneurons, muscles, and glial cells. We validated the efficacy of this toolkit by knocking out known genes in each tissue, demonstrated its orthogonal use with the Gal4/UAS binary expression system, and showed simultaneous knockout of multiple redundant genes. Using these tools, we discovered an essential role for SNARE pathways in NMJ maintenance. Furthermore, we demonstrate that the canonical ESCRT pathway suppresses NMJ bouton growth by downregulating the retrograde Gbb signaling. Lastly, we found that axon termini of motoneurons rely on ESCRT-mediated intra-axonal membrane trafficking to lease extracellular vesicles at the NMJ.

**SIGNIFICANCE STATEMENT:** In this study, we developed a tissue-specific Cas9 toolkit that enables gene knockout specifically in motor neurons, glial cells, and muscle cells, the three cell types of the *Drosophila* peripheral motor system. Complementary to existing RNAi methods, this versatile tissue-specific knockout system offers unique advantages for dissecting gene functions at the neuromuscular junction (NMJ). Using these tools, we discovered that SNARE-mediated secretory pathways are required to maintain the integrity of the NMJ and that ESCRT components play critical yet differential roles in the biogenesis of extracellular vesicles, bouton growth, and membrane turnover at the NMJ. This CRISPR toolkit can be applied to study many biological questions in the neuromuscular system.

## INTRODUCTION

Characterization of developmental mechanisms often involves loss-of-function (LOF) analysis of genes in organisms. Besides the more traditional methods of LOF, such as whole-organismal mutations and RNA interference (RNAi), tissue-specific mutagenesis through the CRISPR/Cas9 system has recently emerged as another powerful approach (1–5). In this approach, either Cas9 or gRNAs (or both) are expressed in a tissue-specific manner, so that CRISPR-mediated mutagenesis of the gene of interest (GOI) occurs only in the desired tissues (1, 2, 6, 7). Although tissue-specific CRISPR can be very effective for studying gene function, its successful application has been limited to a small number of *Drosophila* tissues, including the germline (1, 8), sensory neurons (9–12), prothoracic gland (13), mushroom body neurons (14), imaginal tissues (7, 9, 10), epidermal cells (9, 10), and cardiomyocytes (15). The *Drosophila* neuromuscular junction (NMJ) has been a powerful model for studying many biological processes, such as axon development, synaptogenesis, neuromuscular physiology, and locomotion-related human diseases (16). However, tissue-specific CRISPR has not been successfully applied in it.

NMJs are special synaptic connections formed between motor neurons and somatic muscles (17). The axon termini of the motor neurons release extracellular vesicles (EVs) into the space between axons and muscles (18–21). Outside the nervous system, EVs can originate from many types of cells and can carry diverse cargos, including carbohydrates, lipids, proteins, and nucleic acids (22). These vesicles may function in long-distance signal transduction and cell-cell communication and have been shown to play important roles in many physiological and pathological processes (23, 24). EVs at *Drosophila* NMJs contain several protein and nucleic acid cargos (18, 25) and regulate synaptic structure and activity by mediating axon-to-muscle signal transduction (18, 20). So far, studies of the *Drosophila* NMJ have revealed the requirement of several genes related to intracellular vesicle trafficking and the endocytic pathway in the biogenesis of EVs (19, 21, 24). However, the mechanisms of EV biogenesis at the fly NMJ are not fully understood, and many pathways that are important for EV production in mammalian cells have not been examined in *Drosophila*.

In this study, we generated a tissue-specific CRISPR toolkit for knocking out genes in motor neurons, glia, and somatic muscles, the three cell types of the *Drosophila* NMJ. This toolkit is based on CRISPR-mediated tissue-restricted mutagenesis (CRISPR-TRiM), a method using tissue-specific expression of Cas9 and ubiquitously expressed gRNAs to knock out genes in the desired tissue (9). We validated the efficacy of gene knock-out (KO) in each of the three tissues. Using these tools, we examined the roles of the SNARE pathway and the ESCRT machinery in NMJ morphogenesis. Our results reveal a requirement for the SNARE pathway in NMJ maintenance and critical functions of ESCRT in EV biogenesis, axonal growth, and intra-axonal membrane trafficking.

## RESULTS

### Tissue-specific Cas9 lines for motor neurons, glia, and muscle cells

To apply CRISPR-TRiM in tissues relevant to NMJ biology, we generated several Cas9 lines that are expressed in motoneurons, glial cells, or somatic muscles (Fig. 1*A*) by Gal4-to-Cas9 conversion (10), enhancer-fusion (9), or CRISPR-mediated knock-in (KI). For motoneurons, we generated *wor-Cas9*, which should be active in neuronal progenitor cells (26), and three lines (*OK371-Cas9*, *OK6-Cas9,* and *OK319-Cas9*) that should be expressed in post-mitotic motoneurons (10, 27–30). For glial cells, we previously made *repo-Cas9* (10); for this study, we also generated *gcm-Cas9* by inserting Cas9 into the *gcm* locus through CRISPR-mediate KI (*SI Appendix*, Fig. S1*A*). *gcm-Cas9* is predicted to be active in glial progenitor cells (31, 32). Lastly, we made a muscle-specific *mef2-Cas9* by converting *mef2-Gal4* (10) (Fig. 1*A*).

**Figure 1.**
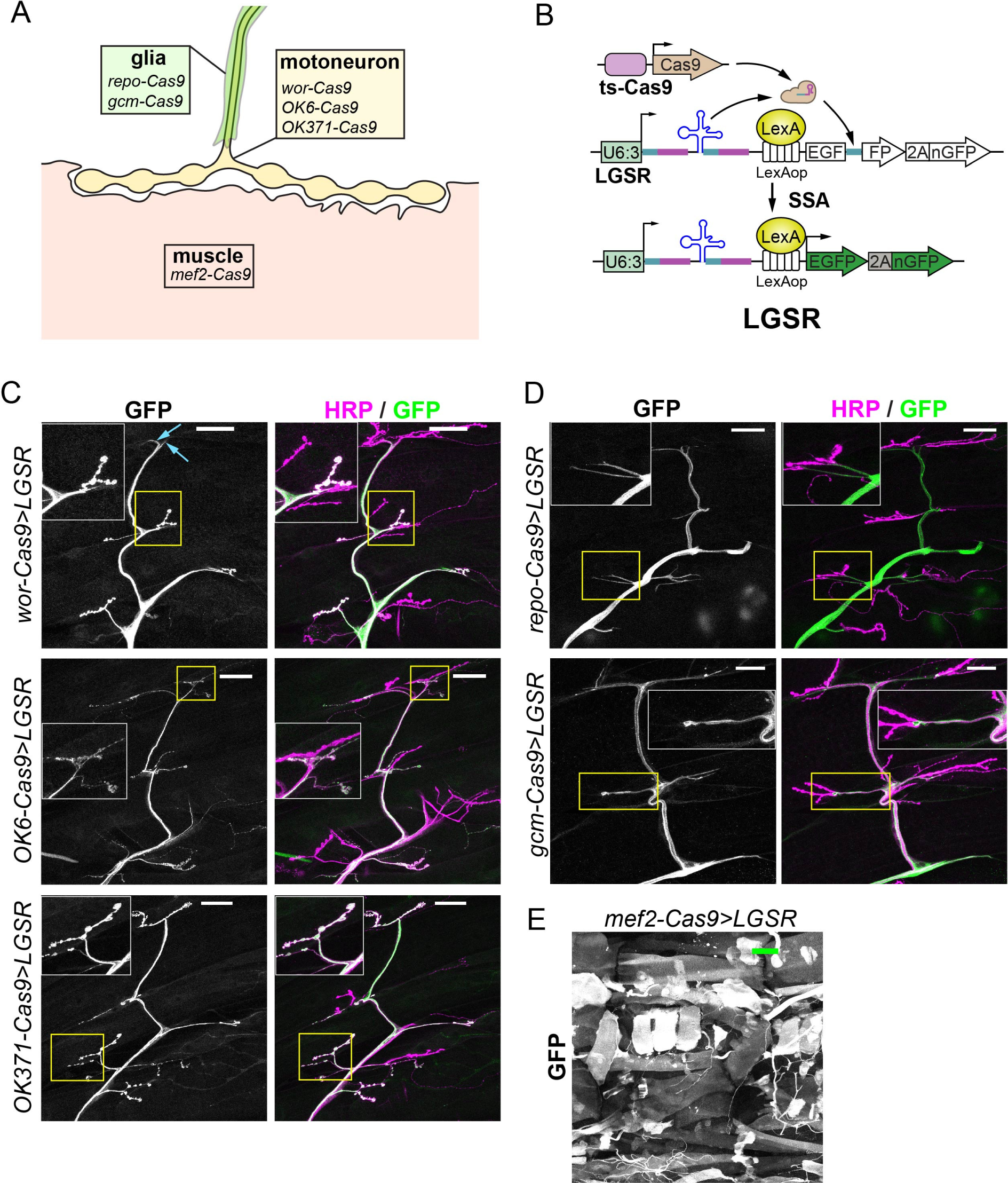
Tissue-specific Cas9 expression patterns characterized by the LGSR reporter. **A**, a diagram of the Cas9 lines made for this study and their targeting tissues. **B**, an illustration of the LGSR reporter. The EGFP coding sequence is reconstituted to give rise to functional protein after gRNA-induced single-strand annealing (SSA). The expression of Cas9 is controlled by a tissue-specific (ts) enhancer. The LexA/LexAop binary system is used to drive strong expression of a cytoplasmic EGFP and a nuclear GFP in cells expressing Cas9. **C**, activity patterns of motoneuron-specific Cas9 lines: *wor-Cas9* (upper panel), *OK6-Cas9* (middle panel) and *OK371-Cas9* (lower panel), characterized with LGSR. Cas9 activity is shown by GFP signal from LGSR reporter. All neurons are shown by HRP antibody immunostaining. Inset: area enclosed by the yellow box showing NMJs. Scale bar: 50μm. **D**, the activity patterns of glia-specific Cas9 lines: *repo-Cas9* (upper panel) and *gcm-Cas9* (lower panel), characterized with LGSR. Neurons are shown by HRP antibody immunostaining. Inset: area enclosed by the yellow box, showing terminal glial protrusions. Scale bar: 50μm. **E**, LGSR labelling of muscle tissues by *mef2-Cas9*. Scale bar: 100μm.

To screen for Cas9 convertants in Gal4-to-Cas9 conversion experiments, we previously made a GFP SSA reporter (GSR), which turns on GFP expression after gRNA-induced single-strand annealing (SSA) and thus can be used to report Cas9 activity (10). Driven by a *ubi* enhancer, GSR signals are relatively weak. To improve the signal intensity, we made LGSR by replacing the *ubi* enhancer in GSR with a *LexAop* enhancer (Fig. 1*B*). By combining a ubiquitous LexA driver (such as *tubP-LexA*), LGSR produces much brighter signals than GSR (*SI Appendix*, Fig. S1*B* and *C*). In use, we found that both GSR and LGSR only label a subset of Cas9-expressing cells, which is reflected by the incomplete labeling of epidermal cells (*SI Appendix*, Fig. S1 *B* and *C*) with the ubiquitous *Act5C-Cas9* (1, 9). This incomplete labeling is likely due to repair of DNA double-strand breaks (DSBs) in GSR by mechanisms other than SSA (33), because in the *lig4* mutant background, where non-homologous end joining (NHEJ) is impaired (34), the same *Act5C-Cas9* resulted in much more complete labeling of cells by GSR (*SI Appendix*, Fig. S1*D*). Thus, GSR and LGSR are useful for visualizing the cell types expressing Cas9 through stochastic labeling but are not faithful reporters of all Cas9-expressing cells.

Using LGSR, we examined the activity patterns of *wor-Cas9*, *OK6-Cas9*, and *OK371-Cas9* at the NMJ (Fig. 1*C*) and in the larval central nervous system (CNS) (*SI Appendix*, Fig. S1*E*), as they should be turned on either early (*wor-Cas9*) or broadly (*OK6-Cas9*, and *OK371-Cas9*) in motoneurons and thus would be useful for CRISPR-TRiM in motoneurons. *wor-Cas9* labeled motoneurons (Fig. 1*C*, upper panel inset) and occasionally glial cells (Fig. 1*C*, upper panel blue arrow) on the larval body wall. In the larval CNS, *wor-Cas9* activated LGSR in many neurons and in some non-neural cells (*SI Appendix*, Fig. S1*E*). These patterns are consistent with the expression of *wor* in neuroblasts (26). At the NMJ, the activities of *OK6-Cas9* and *OK371-Cas9* were mostly restricted to motor neurons (Fig. 1*C*). Unexpectedly, we also observed *OK6-Cas9* activity in many peripheral tissues, including epidermal cells and trachea (*SI Appendix*, Fig. S1*F*). Using the lineage-tracing Gal4 reporter *tubP(FRT.stop)Gal4 UAS-Flp UAS-mCD8::GFP* (10), we confirmed that *OK6-Gal4* is also active in these peripheral tissues during development, in addition to motoneurons (*SI Appendix*, Fig. S1 *G* and *H*). This leaky activity of the *OK6* enhancer in non-neuronal tissues at early developmental stages has been previously reported (35). In the CNS, *OK6-Cas9* showed activity in neurons, as well as some non-neuronal cells, in both optic lobes and the VNC (*SI Appendix*, Fig. S1*E*), while *OK6-Gal4* activity is mostly restricted to the VNC (*SI Appendix*, Fig. S1*I*). Similar to our previous results using the *GSR* reporter (10), we found that LGSR labeled *OK371-Cas9* activity in neurons of the VNC and optic lobes (*SI Appendix*, Fig. S1*E*).

As expected, using LGSR, we detected *repo-Cas9* and *gcm-Cas9* activities in glial cells of the peripheral nerves (Fig. 1*D*) and the larval brain (*SI Appendix*, Fig. S1*J*). Lastly, *Mef2-cas9* showed activity in body wall muscles (Fig. 1*E*), with occasional LGSR activation in epidermal cells and trachea.

In summary, we generated several Cas9 lines that are active in motor neurons, peripheral glia, or somatic muscles and thus are appropriate for tissue-specific mutagenesis at the NMJ.

### Efficient tissue-specific KO of genes in the larval NMJ

To evaluate the efficiency of gene KO in the motor system using CRISPR-TRiM, we generated transgenic gRNAs for several well-studied genes and crossed them to appropriate Cas9 lines. We compared the efficiency of *wor-Cas9*, *OK6-Cas9*, and *OK371-Cas9* in knocking out *Synaptotagmin 1* (*Syt1*) (Fig. 2*A*), which encodes a transmembrane Ca^2+^ sensor on synaptic vesicles (36). *wor-Cas9* resulted in 49.5% reduction of Syt1 protein as assayed by immunostaining (Fig. 2 A *and* D), while *OK6-Cas9* and *OK371-Cas9* caused 74.5% and 89.5% reduction, respectively (Fig. 2*A* and *D*). Considering that biallelic mutagenesis is required to completely knock out a gene, the half reduction of Syt1 in *wor-Cas9* may result from monoallelic mutations in motoneurons. In contrast, *OK6-Cas9* and *OK371-Cas9* should have primarily caused biallelic mutations in *Syt1*.

**Figure 2.**
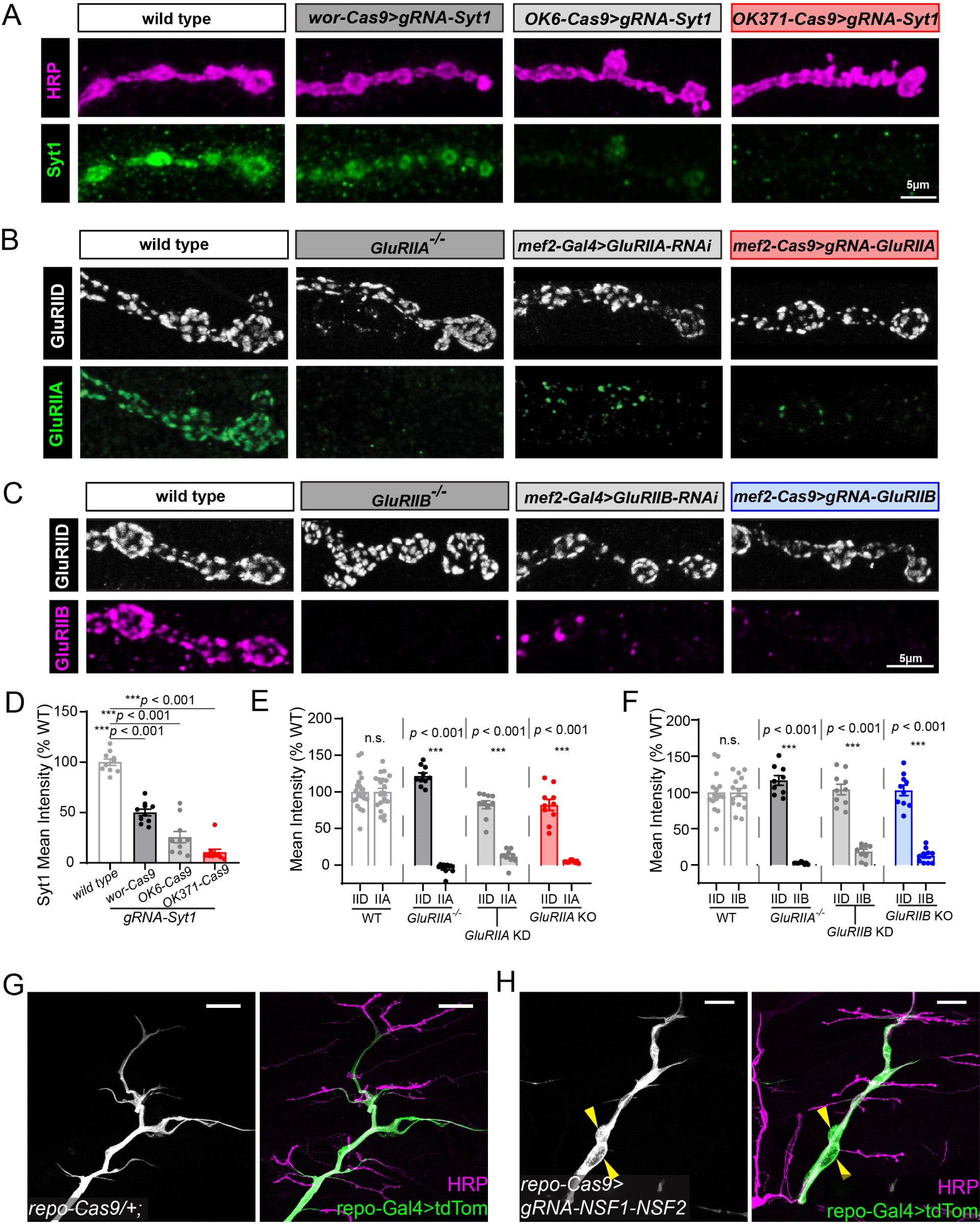
Efficient gene knock out induced by CRISPR-TRiM in the larval motor system. **A**, *Syt1* knock out in motoneurons by *wor-Cas9*, *OK6-Cas9* and *OK371-Cas9*. The Syt1 protein is detected by antibody staining. The axon membrane is shown by HRP staining. Mean intensity of Syt1 signal at NMJ is shown in **D**. **B**, comparison of different methods to remove GluRIIA expression in muscles: whole animal *GluRIIA* mutant (2^nd^ panel), muscle-specific RNAi (3^rd^ panel), and muscle-specific CRISPR KO (4^th^ panel). The GluRIIA protein is detected by antibody staining. GluRIID staining serves as an internal control. Mean intensity of GluRIID and GluRIIA signal at NMJ is shown in **E**. **C**, comparison between different methods to induce *GluRIIB* loss-of-function in muscles. GluRIIB protein level is detected by antibody staining. GluRIID level is unaffected and serves as an internal control. Mean intensity of GluRIID and GluRIIB signal at NMJ is shown in **F**. **D-F**, mean intensities of staining of Syt1 (D), GluRIIA and GluRIID (E), and GluRIIB and GluRIID (F) in the indicated genotypes. One-way ANOVA, p < 0.0001 for all 3 datasets compared to wild type; WT, n=10; wor>Syt1, n=10; OK6>Syt1, n=10; OK371>Syt1, n=10; WT, n=22; GluRIIA^−/−^, n=10; GluRIIA-RNAi, n=10; mef2>GluRIIB, n=10; WT, n=16; GluRIIB^−/−^, n=9; GluRIIB-RNAi, n=10; mef2>GluRIIB, n=10. **G** and **H,** axons and glia in the control (G) and glial-specific KO of *NSF1/NSF2* (H). Glial cells are labeled *repo-Gal4>UAS-CD4-tdTomato* (green), and neurons are labeled by HRP staining (magenta). Yellow arrowheads indicate glial enlargement.

The efficiency of muscle-specific *mef2-Cas9* was evaluated using gRNAs for two *Drosophila* muscle glutamate receptors, GluRIIA and GluRIIB (Fig. 2*B*, *C*, *E* and *F*). In both cases, *mef2-Cas9* efficiently eliminated expression of these proteins in muscle fibers. CRISPR-TRiM generated stronger reduction than RNAi-induced knockdown (KD) using *mef2-Gal4* and was comparable to null mutants of these genes.

To determine the efficiency of *repo-Cas9*, we used gRNAs targeting both *comatose* (*comt or Nsf1*) and *N-ethylmaleimide-sensitive factor 2* (*Nsf2*), which encode two redundant NSF proteins (9) involved in disassembly of SNARE complexes after vesicle fusion (37–39). When paired with a neuronal Cas9, this gRNA transgene results in strong dendrite reduction of *Drosophila* somatosensory neurons (9, 10). As vesicle fusion is an essential “house-keeping” function, we anticipated that loss of both *Nsf* genes in glia should also disrupt glial morphology and function. Indeed, larvae containing both *repo-Cas9* and *gRNA-Nsf1-Nsf2* showed locomotion defects and died at the late third instar stage. In these animals, the glia wrapping motoneuron axons formed enlarged compartments that were absent in *repo-Cas9* controls (Fig. 2 *G* and *H*). These results suggest that *repo-Cas9* can efficiently induce biallelic mutations of two genes simultaneously in glia.

Altogether, the above results demonstrate that Cas9 expressed by motoneurons, muscles, and glial cells can result in efficient tissue-specific gene KO for studying motoneuron development and NMJ biology.

### CRISPR-TRiM reveals a requirement for SNARE components in NMJ maintenance

Having validated the efficacy of CRISPR-TRiM in motoneurons, we next examined the role of the secretory pathway in NMJ morphogenesis. Snap25, Snap24, and Snap29 are *Drosophila* SNARE proteins in the Qbc subgroup that mediate fusion of secretory vesicles with the plasma membrane (40). We have previously generated a transgene expressing six multiplexed gRNAs targeting all three *Snap* genes simultaneously and have showed that it efficiently suppressed dendrite growth of *Drosophila* sensory neurons with an appropriate Cas9 (9). Pairing this gRNA transgene with *wor-Cas9* resulted in apparent locomotion defects since early larval stage and lethality between late 3rd instar and wandering 3^rd^ instar larvae. At 96 h after egg laying (AEL), 41% of the NMJs in these larvae showed morphological defects as compared to the control (Fig. 3*A* and *B*), suggesting that simultaneous KO of all three genes occurred in a mosaic pattern. These NMJs were characterized by large and round boutons with thin connections (Fig. 3*B*) and a 72.1% reduction of bouton numbers (Fig. 3*G*), indicative of NMJ degeneration (41). In addition, large vesicles with strong HRP staining were found in each bouton (*SI Appendix*, Fig. S2 *A* and *B*), suggesting accumulation of intra-axonal membranes. In contrast, when *OK371-Cas9* was used to knock out *Snap* genes, we observed only a moderate reduction of the bouton number across NMJs (Fig. 3 *C*, *D* and *G*). The difference between *wor-Cas9* and *OK371-Cas9* is consistent with our previous results that a Cas9 expressed in neuronal precursor cells is necessary for revealing strong LOF phenotypes of *Snap* genes in somatosensory neurons (9).

**Figure 3.**
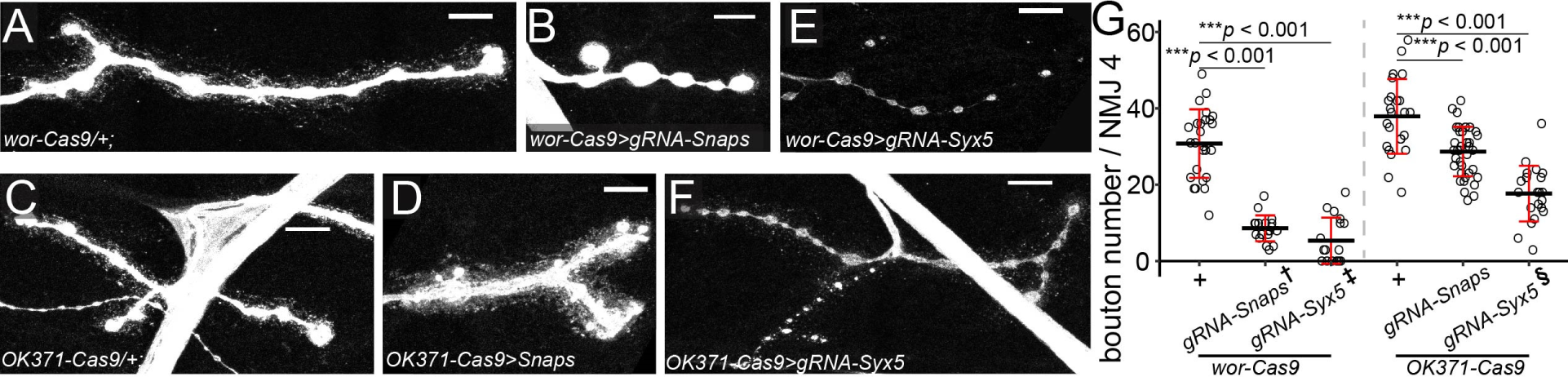
SNARE components are required for NMJ maintenance. **A**–**D,** bouton morphology in *wor-Cas9* (A) and *OK371-Cas9* (C) controls and triple KO of *Snap24*/*Snap25*/*Snap29* in neurons by *wor-Cas9* (B) and *OK371-Cas9* (D). **E** and **F,** bouton morphology resulted from *Syx5* KO in neurons by *wor-Cas9* (E) and *OK371-Cas9* (F). In (A-F), neurons are labeled by HRP staining. Scale bar: 10μm. **G**, bouton numbers in indicated genotypes. One-way ANOVA, F(5,139) = 65.158, *p* < 2.2×10^−16^. Each circle represents an NMJ: *wor-Cas9*, n = 26; *OK371-Cas9*, n= 24; *Snaps^wor-Cas9^*, n = 19; *Snaps^OK371-Cas9^*, n = 38; *Syx5^wor-Cas9^*, n = 17; *Syx5^OK371-Cas9^*, n = 21, between-group *p* values are from multiple comparison test using Bonferroni adjustment. **†**, **‡**, **§** Only the NMJs showing degeneration phenotypes were included in the graph and statistical tests.

*Syx5* encodes a Q-SNARE protein that mediates ER to Golgi transport, an important step in the secretory pathway (Dascher et al., 1994). Using CRISPR-TRiM, we have previously shown that *Syx5* KO in sensory neurons leads to severe dendrite reduction (10). Combining the same *gRNA-Syx5* with either *wor-Cas9* or *OK371-Cas9* resulted in NMJs with several defects, including thinned axons, smaller bouton size, reduced bouton number, and in severe cases, detachment of boutons from axons (Fig. 3*E*, *F* and *G*). The frequency of NMJs showing defects was approximately 40% for both *wor-Cas9* and *OK371-Cas9*, but the phenotype is weaker in *OK371-Cas9* (84.4% bouton reduction) than in *wor-Cas9* (59.8% bouton reduction).

Together, the above data suggest that SNARE-mediated secretory pathways are required for structural maintenance of NMJs and that an early Cas9 is more effective in revealing LOF phenotype of these genes.

### CRISPR-TRiM reveals roles of ESCRT in motoneuron morphogenesis and EV biogenesis

*Drosophila* motoneurons release extracellular vesicles into the synaptic cleft at NMJs. Studies in mammalian cells have uncovered important roles of ESCRT complexes in EV biogenesis (42–44), but whether the ESCRT pathway is involved in EV release at the *Drosophila* NMJ remains unknown. Thus, we investigated the roles of Shrub (Shrb), Tumor susceptibility gene 101 (TSG101) and ALG-2 interacting protein X (ALiX), three components at different steps of the ESCRT pathway, in EV biogenesis at the NMJ using CRISRT-TRiM (Fig. 4*A*).

**Figure 4.**
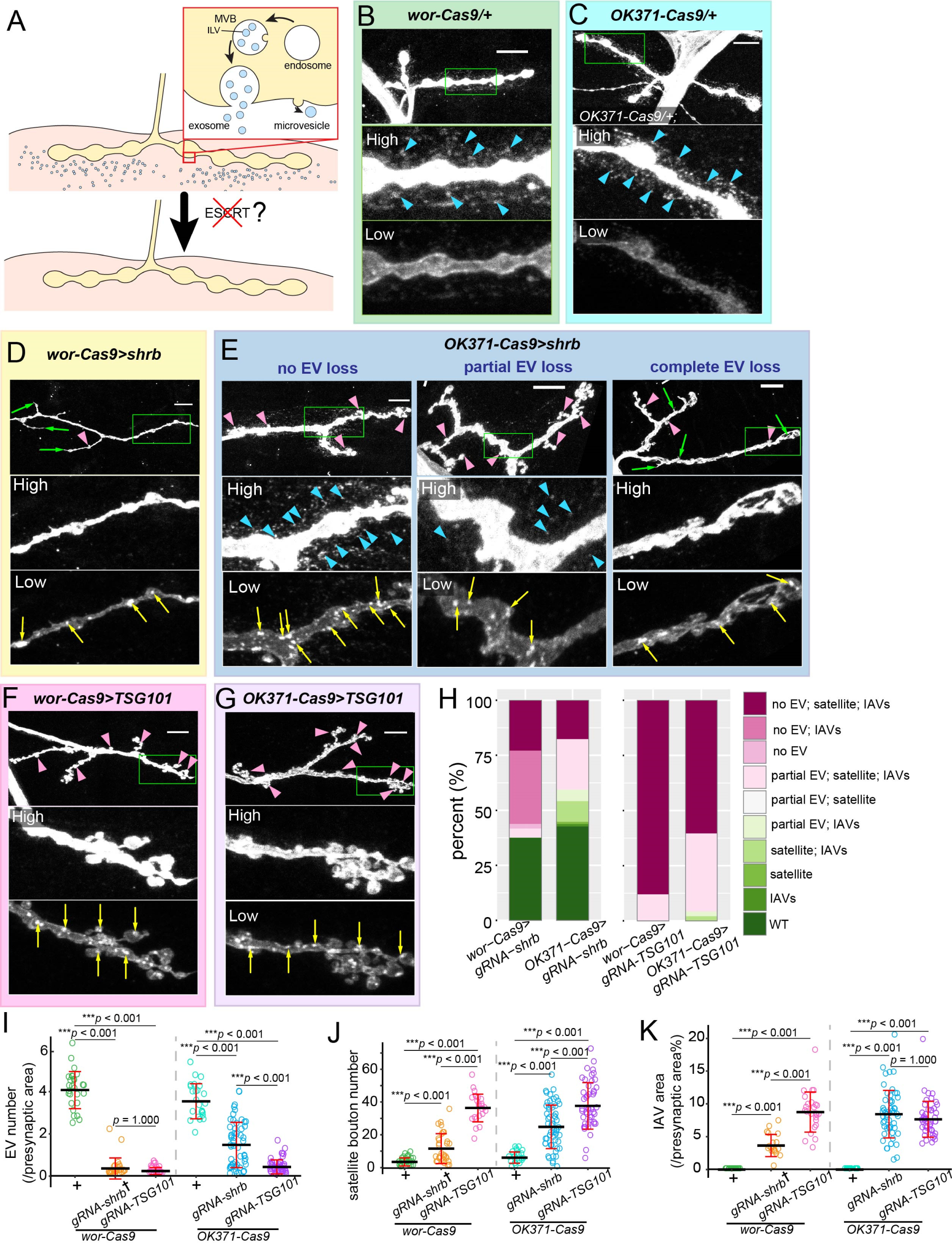
CRISPR-TRiM reveals roles of ESCRT in motoneuron morphogenesis and EV biogenesis. **A**, a diagram of possible routes of EV biogenesis at the NMJ and the experimental design **B**–**G** NMJ morphology in *wor-Cas9* (B) and *OK371-Cas9* (C) controls, *shrb* KO by *wor-Cas9* (D) and *OK371-Cas9* (E), and *TSG101* KO by *wor-Cas9* (F) and *OK371-Cas9* (G). Motoneurons are visualized by HRP staining. “High” and “Low” panels show the zoomed-in views of the area enclosed by the green box imaged with high (to visualize EVs) and low (to visualize IAVs) intensity settings. Three images of *OK371-Cas9>shrb* in (E) show different degrees of EV loss. Blue arrowheads in (B, C, E) indicate the EVs surrounding the presynaptic compartment. Yellow arrows in (D, E, F, G) indicate IAVs. Green arrows in (D and E) indicate filamentous protrusions formed by the presynaptic membrane. Pink arrowheads in (D-G) indicate satellite boutons. Scale bar: 10μm. **H**, percentages of different phenotypes observed in *shrb* and *TSG101* KO experiments. **I**, EV numbers normalized by the presynaptic area. One-way ANOVA, F(5,195) = 146.55, *p* < 2.2×10^−16^. Each circle represents an NMJ: *wor-Cas9*, n = 27; *OK371-Cas9*, n= 24; *shrb^wor-Cas9^*, n = 25; *TSG101^wor-Cas9^*, n = 26; *shrb^OK371-Cas9^*, n = 53; *TSG101^OK371-Cas9^*, n = 46; between-group *p* values are from multiple comparison test using Bonferroni adjustment. **J**, satellite bouton numbers of each NMJ. One-way ANOVA, F(5,203) = 60.643, *p* < 2.2×10^−16^. Each circle represents an NMJ: *wor-Cas9*, n = 26; *OK371-Cas9*, n= 24; *shrb^wor-Cas9^*, n = 30; *TSG101^wor-Cas9^*, n = 25; *shrb^OK371-Cas9^*, n = 57; *TSG101^OK371-Cas9^*, n = 47; between-group *p* values are from multiple comparison test using Bonferroni adjustment. **K**, IAV areas normalized by the presynaptic area. One-way ANOVA, F(5,190) = 73.536, *p* < 2.2×10^−16^. Each circle represents an NMJ: *wor-Cas9*, n = 27; *OK371-Cas9*, n= 24; *shrb^wor-Cas9^*, n = 19; *TSG101^wor-Cas9^*, n = 25; *shrb^OK371-Cas9^*, n = 54; *TSG101^OK371-Cas9^*, n = 47; between-group *p* values are from multiple comparison test using Bonferroni adjustment. **†** Only NMJs with obvious EV loss were included in graphs and statistical tests.

Shrb is the *Drosophila* homolog of Snf7, a central subunit of the ESCRT-III complex, which is responsible for outward budding and fission of vesicles from late endosomes and the plasma membrane (45, 46). We generated *gRNA-shrb* and knocked out *shrb* using *wor-Cas9* and *OK371-Cas9*. In the WT, membranes originated from the axon can be detected by HRP staining (47, 48) including EVs that appear as numerous puncta surrounding axon termini of motoneurons (Fig. 4*B* and *C*, blue arrowheads). *wor-Cas9* together with *gRNA-shrb* resulted in EV loss and morphological changes of axon termini at 63% of NMJs examined (Fig. 4*D* and *H*). Specifically, among all NMJs that displayed any visible phenotypes, 93% lost EVs completely (Fig. 4*D* and *H*). In addition, these NMJs showed 234% increase of satellite boutons (Fig. 4*D*, *H* and *J*, pink arrowheads) and grew filamentous branches (Fig. 4*D*, green arrows). At a lower detection setting, bright HRP-positive puncta (referred to as intra-axonal vesicles or IAVs hereafter, 3.65% presynaptic area) were observed inside distal axons (Fig. 4*D*, yellow arrows). In contrast, IAVs were absent in control neurons (Fig. 4*B* and *K*). Neuroglian (Nrg), a known EV cargo at *Drosophila* larval NMJ (49) (*SI Appendix*, Fig. S3*A*), was accumulated at these IAVs (*SI Appendix*, Fig. S3*B*), indicating mis-trafficking of EV-destined cargos.

When *shrb* was knocked out by *OK371-Cas9*, 57% of NMJs showed phenotypes (Fig. 4*H*). EV levels at these NMJs were variable, ranging from complete loss to almost wildtype levels (Fig. 4*E*, *H* and *K*). In comparison, these NMJs showed consistently strong increases of satellite bouton number (308% increase) and IAV accumulation (8.42% presynaptic area) (Fig. 4*E*, *J* and *K*), suggesting that satellite bouton increase and IAV formation are more sensitive phenotypes of *shrb* LOF than EV loss. As a positive control, we also examined *shrb* KD using *OK371-Gal4*, which resulted in comparable increases in satellite bouton number (383% increase) and IAV accumulation (6.86% presynaptic area) to *OK371-Cas9 gRNA-shrb* but a more complete and consistent EV loss (78.7% reduction) (*SI Appendix*, Fig. S3*C*–*F*). The variation of *OK371-Cas9*-induced KO could be due to variable timings of *shrb* mutations generated in individual neurons. Thus, compared to KD, post-mitotic KO could potentially reveal a broader spectrum of LOF phenotypes.

We next examined TSG101, an ESCRT-I complex component that functions in endosomal cargo sorting and exosome biogenesis (50–52). Unlike *gRNA-shrb*, which caused NMJ defects in only a subset of motoneurons, *gRNA-TSG101* induced near-complete penetrance with both *wor-Cas9* and *OK371-Cas9* (Fig. 4*H*). *wor-Cas9 gRNA-TSG101* showed a complete (100%) penetrance, complete loss of EVs at over 88% of the NMJs examined, a 10-fold increase in satellite bouton number and IAV accumulation (8.73% area) (Fig. 4 F, *I–K*) as compared to the control. *TSG101* KO by *OK371-Cas9* also showed near-complete (98.7%) penetrance (Fig. 4*H*), with much more severe phenotypes of EV loss (87.7% reduction), satellite bouton increase (519% increase), and IAV accumulation (7.64% area) (Fig. 4 *G*, *I–K*) compared to those of *OK371-Cas9 gRNA-Shrb*. Thus, for both Cas9s, *TSG101* KO produced strongest and similar levels of EV loss, satellite bouton increase, and IAV accumulation, suggesting that both pre-mitotic and post-mitotic KO of *TSG101* may reveal the null phenotype.

Lastly, we knocked out *ALiX*, which encodes a BRO1 domain-containing protein that can recruit the ESCRT-III complex, as ALiX has been shown to be involved in EV biogenesis in mammalian cells (44, 53, 54). ALiX functions upstream of ESCRT-III in an alternative pathway to the canonical pathway mediated by ESCRT-0, I, and II complexes (53–55). We generated *gRNA-ALiX* and validated its efficiency using the Cas9-LEThAL assay (9). However, even pre-mitotic *ALiX* KO by *wor-Cas9* did not exhibit any noticeable morphological defects (*SI Appendix*, Fig. S4*G–I*), suggesting that motor neurons use the canonical ESCRT pathway, not the ALiX-assisted pathway, to generate EVs.

Together, the above data suggest that the canonical ESCRT pathway is required for both EV biogenesis and for suppressing the growth of satellite boutons. However, the severities of these two phenotypes do not always correlate, suggesting that they may be controlled by divergent pathways downstream of TSG101 and Shrb.

### ESCRT LOF causes satellite bouton overgrowth by upregulating BMP signaling

The growth of satellite boutons at the *Drosophila* NMJ relies on the muscle-derived BMP ligand Glass-bottom boat (Gbb), which binds to presynaptic BMP receptors. The BMP receptors on the axon membrane are then downregulated by endocytosis (56, 57). Defects of endocytosis in neurons result in excessive satellite boutons due to persistent ligand-receptor interaction (58, 59). Because ESCRT-mediated cargo sorting is required for terminating ligand/receptor signaling (50, 60–62), ESCRT could suppress satellite bouton growth by downregulating Gbb signaling. Alternatively, it may do so by generally promoting material turnover at the NMJ through the lysosomal pathway. To distinguish these two possibilities, we asked whether the satellite bouton increase associated with ESCRT LOF depends on Gbb. Consistent with published results (56), global KD of *gbb* by *Act-Gal4* (Fig. 5*B*) caused 80.7% reduction of satellite boutons (Fig. 5*G*) and 23.2% reduction of axial boutons (Fig. 5*H*) as compared to the control. Strikingly, *gbb* KD reduced satellite boutons in *TSG101* KO and *shrb* KO (by *wor-Cas9*) to the control level (Fig. 5 *D*, *F* and *G*), suggesting that Gbb signaling is responsible for the increase of satellite boutons in ESCRT LOF. Interestingly, besides excess satellite boutons, *TSG101* KO also resulted in 18.2% increase of axial boutons (Fig. 5*H*), but this increase was not affected by *gbb* KD (Fig. 5*F*). We did not observe obvious axial bouton increase in *shrb* KO (Fig. 5*H*). These data suggest that TSG101, but not Shrb, also has a Gbb-independent role in suppressing NMJ overgrowth. Lastly, the filamentous protrusions associated with *shrb* KO were still present after *gbb* KD (Fig. 5*F*), suggesting that these structures are independent of Gbb signaling.

**Figure 5.**
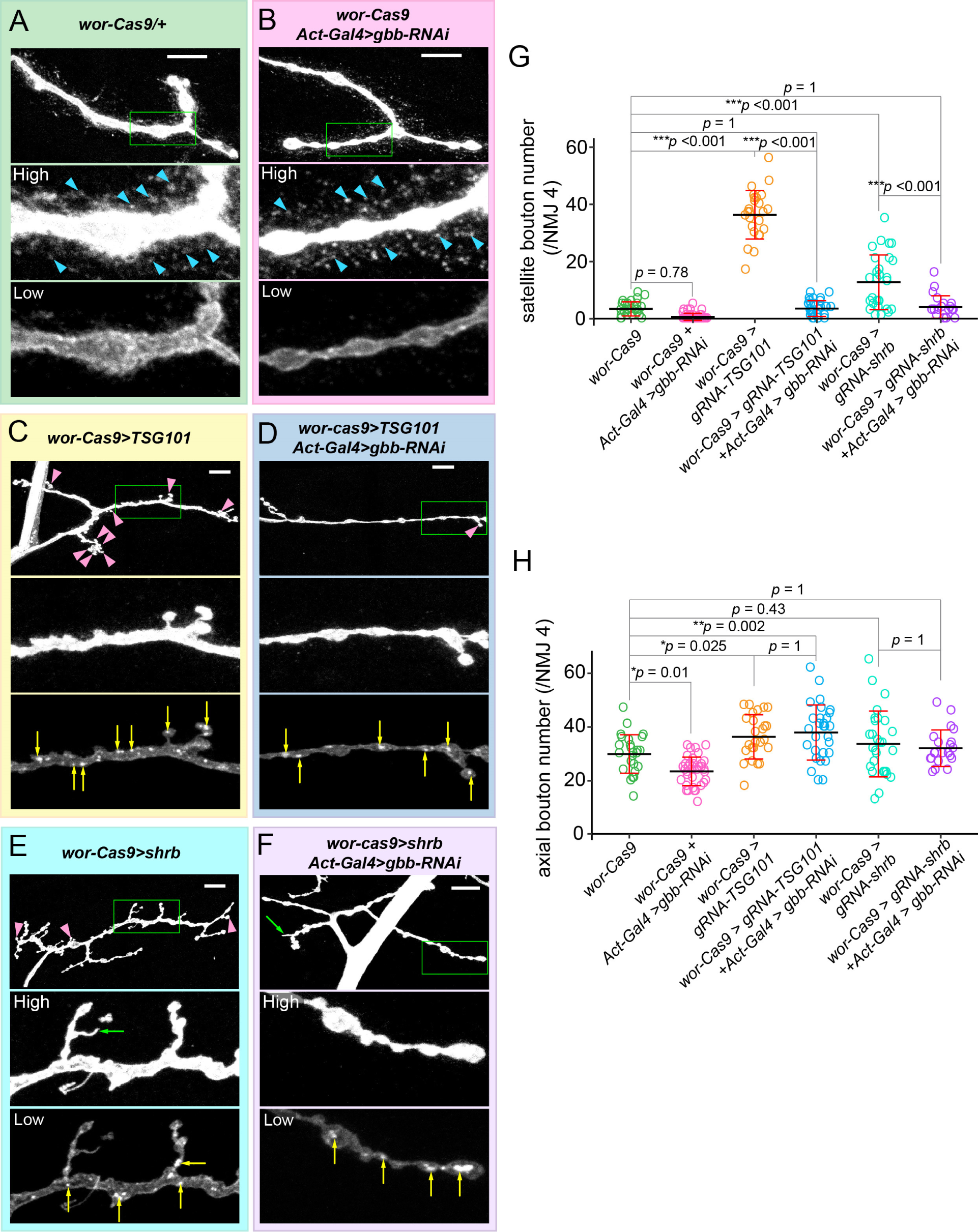
ESCRT LOF causes satellite bouton overgrowth by upregulating BMP signaling. **A**–**F,** NMJ morphologies in the control (A), global *gbb* KD (B), *TSG101* KO by *wor-Cas9* (C), *TSG101* KO combined with *gbb* KD (D), *shrb* KO by *wor-Cas9* (E) and *shrb* KO combined with *gbb* KD (F). Motoneurons are visualized by HRP staining. “High” and “Low” panels show the zoomed-in views of the area enclosed by the green box. The same NMJ is imaged with both high and low intensity settings. Pink arrowheads indicate satellite boutons. Yellow arrows indicate IAVs. Blue arrowheads indicate EVs. Green arrowheads indicate protrusions from axons. Scale bar: 10μm. **E**, satellite bouton numbers in the indicated genotypes. One-way ANOVA, F(5,159) = 153.58, *p* < 2.2×10^−16^. Each circle represents an NMJ: *wor-Cas9*, n= 26; *gbb-RNAi^Act-Gal4^*, n = 36; *TSG101^wor-Cas9^*, n = 25; *TSG101^wor-Cas9^ / gbb-RNAi^Act-Gal4^*, n = 29; *shrb^wor-Cas9^*, n = 28; *shrb^wor-Cas9^ / gbb-RNAi^Act-Gal4^*, n = 21; between-group *p* values are from multiple comparison test using Bonferroni adjustment. **F**, axial bouton numbers in indicated genotypes. One-way ANOVA, F(5,159) = 11.555, *p* = 1.57×10^−9^. Each circle represents an NMJ: *wor-Cas9*, n= 26; *gbb-RNAi^Act-Gal4^*, n = 36; *TSG101^wor-Cas9^*, n = 25; *TSG101^wor-Cas9^ / gbb-RNAi^Act-Gal4^*, n = 29; *shrb^wor-Cas9^*, n = 28; *shrb^wor-Cas9^ / gbb-RNAi^Act-Gal4^*, n = 21; between-group *p* values are from multiple comparison test using Bonferroni adjustment. The datasets of *wor-Cas9>gRNA-shrb* and *wor-Cas9>gRNA-TSG101* in (G) and (H) are the same as in Figure 4.

To further understand whether the roles of TSG101 and Shrb in EV biogenesis and intra-axonal membrane turnover are related to Gbb signaling, we also examined EV and IAV levels in *TSG101* and *shrb* KO combined with *gbb* KD. Global *gbb* KD did not affect the EV level by itself (Fig. 5*B* and *SI Appendix*, Fig.S4*A*), nor did it change the EV level in *TSG101* or *shrb* KO (Fig. 5 *D* and *F*, and *SI Appendix*, Fig.S4*A*), confirming that EV biogenesis and satellite bouton growth are controlled by separate pathways downstream of ESCRT. Interestingly, *gbb* KD caused a 40.0% reduction of IAVs in *TSG101* KO but did not significantly change IAV levels in *shrb* KO (*SI Appendix*, Fig.S4*B*). These results suggest that IAV accumulation in *TSG101* KO is partially due to Gbb signaling.

## DISCUSSION

### CRISPR-TRiM is a versatile tool for dissecting gene function in the motor system

Although tissue-specific mutagenesis by CRISPR is a powerful approach for dissecting gene functions in animal development (1, 4, 9–12, 14, 63), its successful application in the *Drosophila* NMJ system has not been demonstrated previously. In this study, we developed a Cas9 collection for applying CRISPR-TRiM in motor neurons, somatic muscles, and glia cells, the three principal cell types that make up the NMJ. Using these tools, we demonstrate the effectiveness of gene KO in each tissue and reveal the role of the SNARE pathway in NMJ maintenance and the roles of the ESCRT pathway in NMJ morphogenesis.

Compared to LOF by RNAi, the CRISPR-TRiM method offers two major advantages. First, it can efficiently mutagenize multiple genes simultaneously and thus is particularly useful for studying redundant genes, as exemplified by the analyses of *Nsf* and *Snap* genes. Second, because mutagenesis occurs individually in each cell and could yield different degrees of LOF, CRISPR-TRiM has the potential to reveal both weak and strong LOF phenotypes of the gene of interest (GOI) using a single gRNA transgene. This property can be useful for dissecting complex functions of genes, as demonstrated by *OK371-Cas9 gRNA-shrb*. Unlike Gal4-dependent CRISPR strategies (1, 7), the tissue-specific Cas9s in our CRISPR-TRiM method can be used orthogonally with binary expression systems, as exemplified by simultaneous *TSG101* KO in neurons and global KD of *gbb*. Thus, the tools we reported here enable a versatile CRISPR toolkit to analyze gene function in the NMJ system.

### Factors determining the efficacy of CRISPR-TRiM

To reveal the null phenotype of a gene at the single cell level, biallelic LOF mutations need to be generated early in the cell’s lineage (9). In practice, multiple factors can influence the timing and the nature of mutations, and thus the efficacy of tissue-specific KO.

First, the expression timing, duration, and strength of the Cas9 can largely affect the extent of LOF. Cas9s that are expressed in neural stem cells or progenitor cells can result in earlier mutations than those that are expressed in postmitotic neurons. An example is that, with the same *gRNA-shrb*, *wor-Cas9* caused more severe EV loss than *OK371-Cas9*. On the other hand, post-mitotic Cas9s presumably have stronger and more long-lasting expression than precursor-cell Cas9s. This property could be important for mutagenesis with gRNAs that have slow kinetics. *gRNA-Syt1* may be an example of slow gRNAs that require the sustained activity of *OK371-Cas9* to induce biallelic mutations.

Second, the efficiency of gRNAs critically affects the outcome of KO. The gRNA efficiency is affected by not only the target sequence (64, 65), the construct design (6, 7, 66, 67), the accessibility of the target sequence (65, 68–70), but also the functional significance of the mutated amino acids. If in-frame mutations at the target site do not completely disrupt a protein’s function, a portion of the cells carrying even biallelic mutations will display no or hypomorphic phenotypes. This scenario could have contributed to the motor neurons that showed no defects in *shrb* and *Syx5* KO and possibly the variable phenotypes of *OK371-Cas9 gRNA-shrb* neurons. However, variable phenotypes for the same gene can be advantageous for revealing multiple facets of a gene’s function, as in this case.

Lastly, the expression timing and product stability of the GOI can affect the severity of the LOF phenotype. For genes that are expected to express late in the cell lineage, such as only in differentiating neurons, post-mitotic Cas9 can be early enough for causing LOF (9). *Syt1* may fall in this category. In contrast, house-keeping genes are typically expressed earlier than tissue-specific Cas9s and are thus more prone to perdurance effects (9). For these genes, early expressing Cas9s should be more effective than late Cas9s, as in the case of the KO for *shrb*, *Syx5*, and *Snap* genes. However, if the mRNA/protein products of the GOI are rapidly turned over, even house-keeping genes could be effectively removed by a late expressing Cas9. This scenario may explain *TSG101* KO results.

In summary, the efficacy of CRISPR-TRiM is influenced by Cas9 expression pattern, gRNA efficiency, and the characteristics of the GOI. In conducting CRISPR-TRiM, optimal results can be achieved by choosing the appropriate combinations of Cas9 and gRNAs. Interpretation of the results should also take consideration of the property of the GOI.

### The ESCRT pathway controls multiple aspects of NMJ morphogenesis

By examining *shrb* and *TSG101* KO, we uncovered several aspects of NMJ morphogenesis controlled by the ESCRT pathway, namely EV biogenesis, satellite bouton growth, and intra-axonal membrane trafficking. Specifically, disruptions of the ESCRT pathway resulted in EV loss, overgrowth of satellite boutons, and accumulation of IAVs. Our results suggest that these phenotypes are controlled by both shared and separate pathways downstream of ESCRT components.

First, the ESCRT pathway suppresses satellite bouton growth by downregulating Gbb signaling. This function of ESCRT is consistent with its role in sorting signaling receptors to intralumenal vesicles (ILVs) of the multivesicular body (MVB) for subsequent degradation in lysosomes (71). In the absence of ESCRT, MBP receptors may remain associated with Gbb or be recycled back to the axon membrane to sustain the signaling.

Second, we found that the ESCRT pathway is essential for EV biogenesis at the NMJ. The phenotypic spectrum of *shrb* KO suggests that EV loss and satellite bouton overgrowth are two uncorrelated defects. Global *gbb* KD in ESCRT LOF further confirmed that the EV loss and satellite bouton overgrowth are controlled by two separate pathways. The importance of ESCRT in EV biogenesis has been reported in other systems previously (22, 43, 44, 52). EVs are generated through either fusion of MVBs with the plasma membrane (exosomes) or outward budding of vesicles from the plasma membrane (microvesicles) (22), two processes that both require the ESCRT machinery (72). However, the roles of ESCRT in the biogenesis of axon-derived EVs have not been characterized previously. Our results confirmed that the EVs at the NMJ also depend on the ESCRT.

Third, we found that ESCRT prevents accumulation of IAVs by both Gbb-dependent and independent mechanisms. ESCRT is known to be important for endomembrane turnover by generating ILVs that are subsequently degraded in the lysosome (73). For this reason, disruptions of ESCRT could cause accumulation of late endosomes and give rise to IAVs. The enrichment of the anti-HRP epitope suggests that these vesicles may normally feed into degradative compartments. Interestingly, the elevated level of Gbb signaling caused by *TSG101* KO exacerbates this phenotype, possibly by stimulating biogenesis and delivery of membranes to axons termini and further clogging the system.

In addition to these three main phenotypes, we also observed a mild increase of axial bouton growth in *TSG101* KO, which is independent of the Gbb signaling. This phenomenon could be related to TSG101’s role in material turnover at the NMJ. Because ESCRT is upstream of EV release and lysosomal degradation, both of which effectively result in membrane disposal, disrupting ESCRT could pile up membranes at the NMJ and contribute to bouton expansion.

Lastly, our data provide interesting clues about how EVs are generated by axons. We found that the EV cargo Nrg is accumulated at IAVs in *shrb* KO, suggesting that the EV cargo is sorted to these endosomal compartments before release. Thus, it appears that at least some EVs are exosomes generated through the MVB pathway. However, this observation does not rule out the possibility of EV biogenesis through microvesicle budding at the plasma membrane.

### Shared and separate pathways downstream of ESCRT components at the NMJ

Our analyses of three ESCRT components show that their LOF does not produce identical phenotypes. First, while mutant neurons of *shrb* and *TSG101* both show near complete EV loss, *shrb* mutant neurons with extreme morphological defects did not exhibit as strong increases of satellite and axial boutons and IAV accumulation as *TSG101* mutant neurons. Instead, they grew filamentous membrane protrusions, which were absent in *TSG101* mutant neurons. In addition, Gbb signaling contributes to IAV accumulation in *TSG101* KO but not in *shrb* KO. Thus, the Gbb signaling seems to contribute to the NMJ defects of *TSG101* KO much more strongly than to those of *shrb* KO. These differences could be due to TSG101 and Shrb functioning at different steps of signaling receptor processing on endosomal membranes and/or that ESCRT-III is required for more molecular processes than those involving ESCRT-I (46, 74–76). Second, we found that *ALiX* KO does not affect NMJ morphology. ALiX acts in parallel with ESCRT-I to direct ubiquitinated cargo to ESCRT-III (53). ALiX has also been shown to be involved in EV biogenesis (53, 54). However, we found that ALiX does not contribute to EV biogenesis at the NMJ, suggesting that ALiX’s role in EV formation may be context-dependent.

## MATERIALS AND METHODS

### *Drosophila* stocks and culture

The details of fly strains used in this study are listed in Key Resource Table. All crosses were set up in the standard yeast-sugar fly food and kept at 25℃ and 60% humidity, with 12 h light/dark cycle until examination.

### Molecular cloning

*LGSR*: A DNA fragment containing 13x LexAop2 enhancer, hsp70 core promotor, and a synthetic intron was PCR-amplified from pDEST-APLO (Addgene 112805) (77). a GFP single-strand annealing cassette, which contains the following sequences in order: EGFP coding sequence for amino acid 1 to amino acid 175, a synthetic fragment (CCGTCTGTCACAGGATTGGCTGCTTG) that serves as gRNA targeting sites, and EGFP coding sequence for amino acid 43 to amino acid 233, was PCR-amplified from pAC-GSR (10). Both fragments were assembled into MluI/BglII sites of pAC-GSR through NEBuilder HiFi DNA assembly (New England Biolabs, Inc) to make pAC-LGSR.

*gcm-Cas9*: Two gRNA spacer sequence targeting the 5’UTR immediately before the start codon of gcm were first cloned into pAC-U63-tgRNA-nlsBFPnls (Addgene 169029) (78) according to published protocols. The resulting plasmid was digested by PstI and assembled with three DNA fragments through NEBuilder HiFi DNA assembly to make a *gcm* gRNA-donor vector. The three DNA fragments are a 5’ homology arm (827 bp immediately before *gcm* start codon), in which the gRNA target sequences were mutated, a Cas9-T2A fragment PCR amplified from pDEST-APIC-Cas9 (Addgene 121657) (9), and a 3’ homology arm (966 bp immediately after the start codon).

*gRNA transgenic vectors*: gRNA target sequences for genes of interest were cloned into various gRNA vectors according to published protocols (9, 10). The gRNA target sequences are listed in Table 1, and the cloning vectors and the PCR templates are listed in Table 2.

**Table 1.**
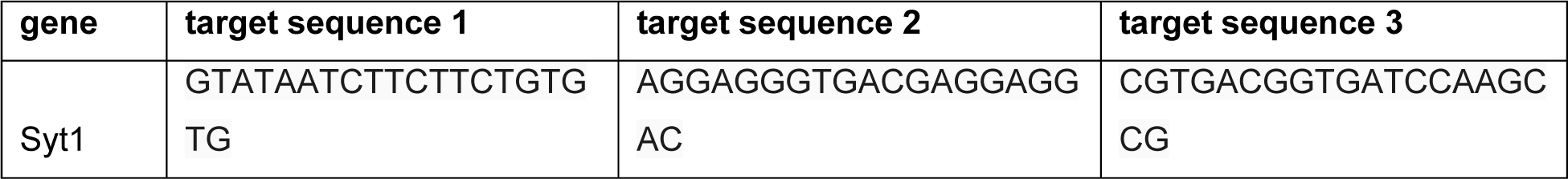
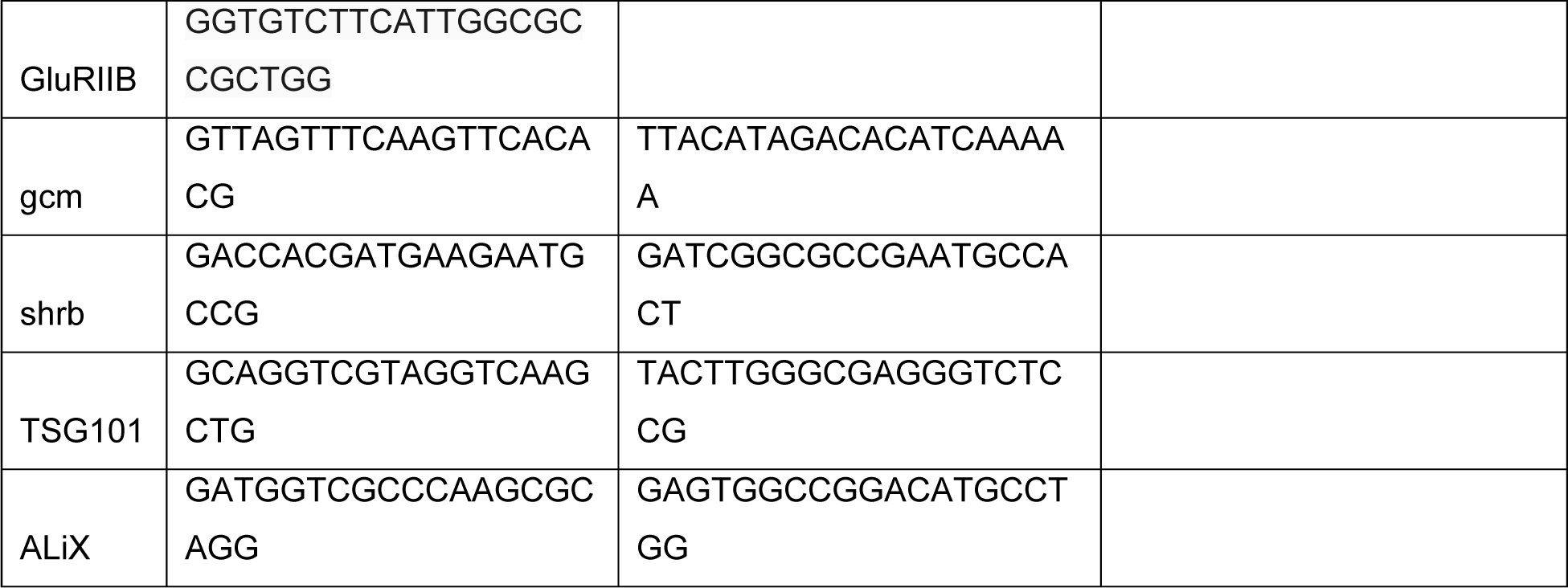
gRNA target sequences.

**Table 2.** gRNA expression vectors.

| gRNA lines | gRNA cloning vector | PCR template |
| --- | --- | --- |
| gRNA-Syt1 | pAC-U63-tgRNA-Rev | pMGC |
| gRNA-GluRIIB | pAC-U63-tgRNA-Rev | NA |
| gRNA-shrb | pAC-U63-tgRNA-Rev | pMGC |
| gRNA-TSG101 | pAC-U63-tgRNA-Rev | pMGC |
| gRNA-ALiX | pAC-U63-tgRNA-Rev | pMGC |

Transgenic constructs were injected by Rainbow Transgenic Flies (Camarillo, CA, USA) to transform flies through φC31 integrase-mediated integration into attP docker sites.

### Generation of tissue-specific Cas9 lines

*wor-Cas9*, *OK6-Cas9*, and *mef2-Cas9* were converted from corresponding Gal4 lines using the HACK method as previously described (10). GSR was used as the reporter for Cas9 conversion. A 2nd chromosomal donor was used to convert 2nd chromosomal Cas9 (*wor-Cas9* and *OK6-Cas9*) and a 3rd chromosomal donor was used to convert 3rd chromosomal Cas9 (*mef2-Cas9*).

To make *gcm-Cas9*, the *gcm* gRNA-donor vector was first inserted into the *VK3a* attP site (79) by φC31 integrase-mediated integration. This gRNA-donor transgene was then crossed to *y[1] nos-Cas9.P[ZH-2A] w[*]* (1) to induce homologous recombination in the germline of the progeny. The resulting male progeny were crossed to the Cas9 positive tester *Act-Gal4 UAS-EGFP; tub-Gal80 gRNA-Gal80* (9) for screening larvae that showed GFP signals in the brain. The larvae were recovered and used to derive isogenic *gcm-Cas9* strains by removing transgenic components on other chromosomes. 5 larvae showing identical GFP patterns in the brain were recovered from 172 larvae in total.

### Validation of gRNA efficiency

The efficiency of transgenic gRNA lines was validated by the Cas9-LEThAL assay (9). Homozygous males of each gRNA line were crossed to *Act-Cas9 w lig4* homozygous females. *gRNA-Syt1, gRNA-GluRIIA, and gRNA-GluRIIB* crosses resulted in lethality before pupation; *gRNA-shrb* crosses resulted in lethality of all progeny in embryos; *gRNA-TSG101* crosses resulted in lethality of all progeny in 1^st^ to 2^nd^ instar larvae; *gRNA-ALiX* crosses resulted in lethality of male progeny from 3^rd^ instar larvae to pharate adults and viable and healthy female progeny. These results suggest that all gRNAs are efficient.

### Live Imaging

Live imaging was performed as previously described (77). Briefly, animals were reared at 25°C in density-controlled vials for 120 hours (wandering third instar). Larvae were mounted in glycerol and imaged using a Leica SP8 confocal microscope.

### Larval fillet preparation

Larval fillet dissection was performed on a petri dish half-filled with PMDS gel. Wandering third instar larvae were pinned on the dish in PBS dorsal-side up and then dissected to expand the body wall. PBS was then removed and 4% formaldehyde in PBS was added to fix larvae for 15 minutes at room temperature, or Bouin’s solution was added for 5 minutes at room temperature. Fillets were rinsed and then washed at room temperature in PBS for 20 minutes or until the yellow color from Bouin’s solution faded. After immunostaining, the head and tail of fillets were removed, and the remaining fillets were placed in SlowFade Diamond Antifade Mountant (Thermo Fisher Scientific) on a glass slide. A coverslip was lightly pressed on top. Larval fillets were imaged with 40× or 63× oil objectives using a Leica SP8 confocal microscope.

### Larval brain preparation

Larval brain dissection was performed as described previously (80). Briefly, wandering 3rd instar larvae were dissected in a small petri dish filled with cold PBS. The anterior half of the larva was inverted, and the trachea and gut were removed. Samples were then transferred to 4% formaldehyde in PBS and fixed for 25 minutes at room temperature. Brain samples were washed with PBS. After immunostaining, the brains were placed in SlowFade Diamond Antifade Mountant on a glass slide. A coverslip was lightly pressed on top. Brains were imaged with both 20× and 40× oil objectives using a Leica SP8 confocal microscope.

### Immunohistochemistry

For larval brains: Following fixation, brains were rinsed and then washed twice at room temperature in PBS with 0.3% Triton-X100 (PBST) for 20 minutes each. Brains were then blocked in a solution of 5% normal donkey serum (NDS) in PBST for 1 hour. Brains were then incubated in the blocking solution with rat mAb 7E8A10 anti-elav (1:10 dilution, DSHB) or mouse mAb 8D12 anti-repo (1:20 dilution, DSHB) overnight at 4°C. Following incubation brains were then rinsed and washed in PBST 3 times for 20 minutes each. Brains were then incubated in a block solution containing a donkey anti-rat or donkey anti-mouse secondary antibody conjugated with Cy5 or Cy3 (1:400 dilution, Jackson ImmunoResearch) for 2 hours at room temperature. Brains were then rinsed and washed in PBST 3 times for 20 minutes each and stored at 4°C until mounting and imaging.

For larval fillets: following fixation, fillets were rinsed and then washed at room temperature in PBS. Fillets were then removed from PMDS gel and blocked in a solution of 5% normal donkey serum (NDS) in 0.2% PBST for 1 hour. Fillets were then incubated in the blocking solution with primary antibodies overnight at 4°C. Primary antibodies used in this study are mouse mAb BP 104 anti-Neuroglian (1:8 dilution, DSHB), mouse mAb 8B4D2 anti-GluRIIA (1:50, DSHB), rabbit anti-Syt (1:2500); (81), rabbit anti-GluRIIB (1:1000) (41), rabbit anti-vGluT (1:200 dilution, generated using the same peptide and approach described in (82)), guinea pig anti-GluRIID (1:1000) (41), and mouse mAb 4F3 anti-discs large (1:20 dilution, DSHB). Following incubation brains were then rinsed and washed in PBST 3 times for 20 minutes each. Brains were then incubated in a block solution containing fluorophore-conjugated conjugated secondary antibodies for 2 hours at room temperature. Secondary antibodies used in this study are: goat anti-HRP conjugated with Cy3 (1:200, Jackson ImmunoResearch), donkey anti-mouse secondary antibody conjugated with Cy5 or Alexa 488 (1:400, Jackson ImmunoResearch), donkey anti-rabbit secondary antibody conjugated with Cy5 or Alexa488 (1:400, Jackson ImmunoResearch) and rabbit polyclonal anti-GFP antibody conjugated with Alexa 488 (1:400, LifeTechnologies, A21311). Fillets were then rinsed and washed in PBST 3 times for 20 minutes each and stored at 4°C until mounting and imaging.

### Image analysis and quantification

Muscle 4 NMJ of segment A2‒A6 was used for quantification. ImageJ software was used to quantify EV numbers in batches. Images were segmented by the Trainable Weka Segmentation plugin. HRP antibody staining results were used to quantify the EV number and IAV coverage. The machine learning-based program was first trained by several sample images to distinguish the background, presynaptic compartment, and EV/IAV. Then the model was applied to segment all the images in control and experimental groups of the same experiment. The EV number and IAV coverage were measured using Analyze Particles function in ImageJ.

The EV number at each NMJ is normalized. The total EV number in each region of interest (ROI) was divided by the presynaptic membrane area. The IAV coverage (%) was also normalized by dividing the area of IAV by the corresponding presynaptic membrane area.

The numbers of axial and satellite boutons were manually counted, according to the imaging results from vGluT and Dlg staining.

### Experimental design and statistical analysis

For all experiments, the control groups and the experimental groups were kept in the same growing conditions. The same dissection and staining procedures were applied to all the groups. The animals used for dissection were of the same age (∼120h AEL wandering 3rd instar larva). RStudio was used to perform one-way analysis of variance (ANOVA) and Student’s t-test where indicated. For experiments involving only two groups, a two-tailed t-test was used to compare the means. Non-equal variance was assumed. For experiments with more than two groups, one-way ANOVA was first applied to identify significantly different mean(s). After that, multiple comparisons were performed using the Bonferroni post hoc method.

## Supporting information

Resource table

## DATA SHARING PLANS

All study data are included in the article and/or Supplemental Methods. Constructs and fly strains will be available from Addgene and Bloomington *Drosophila* Stock Center, respectively, or upon request.

## ACKNOWLEDGEMENT

We thank the Bloomington *Drosophila* Stock Center (BDSC) for fly stocks; Claire Amber Ho for technical help; Avital Rodal and Erica Dresselhaus (Braindeis University) for discussion and communication about unpublished results; Ellie Heckscher (University of Chicago) for suggestions of motoneuron precursor Gal4; Scott Emr and Inle Bush for reading and suggestions on this manuscript. This work was supported by NIH grants (R01NS099125, R21OD023824, and R24OD031953) awarded to C.H., and by NIH grants (R01NS126654) awarded to D.D..

## Author contributions

Han C and Chen X designed the experiments; Wang B conducted molecular cloning; Chen X and Hu J performed dissection, immunostaining, and imaging; Wang S and Chen X developed the quantification methods; Chen X performed quantification; Han C, Chen X, and Perry S and Loxterkamp E built genetic reagents used in this study; Han C and Chen X wrote the manuscript; Han C, Chen X, Dickman D, and Perry S revised the manuscript.

---

**Figure S1.**
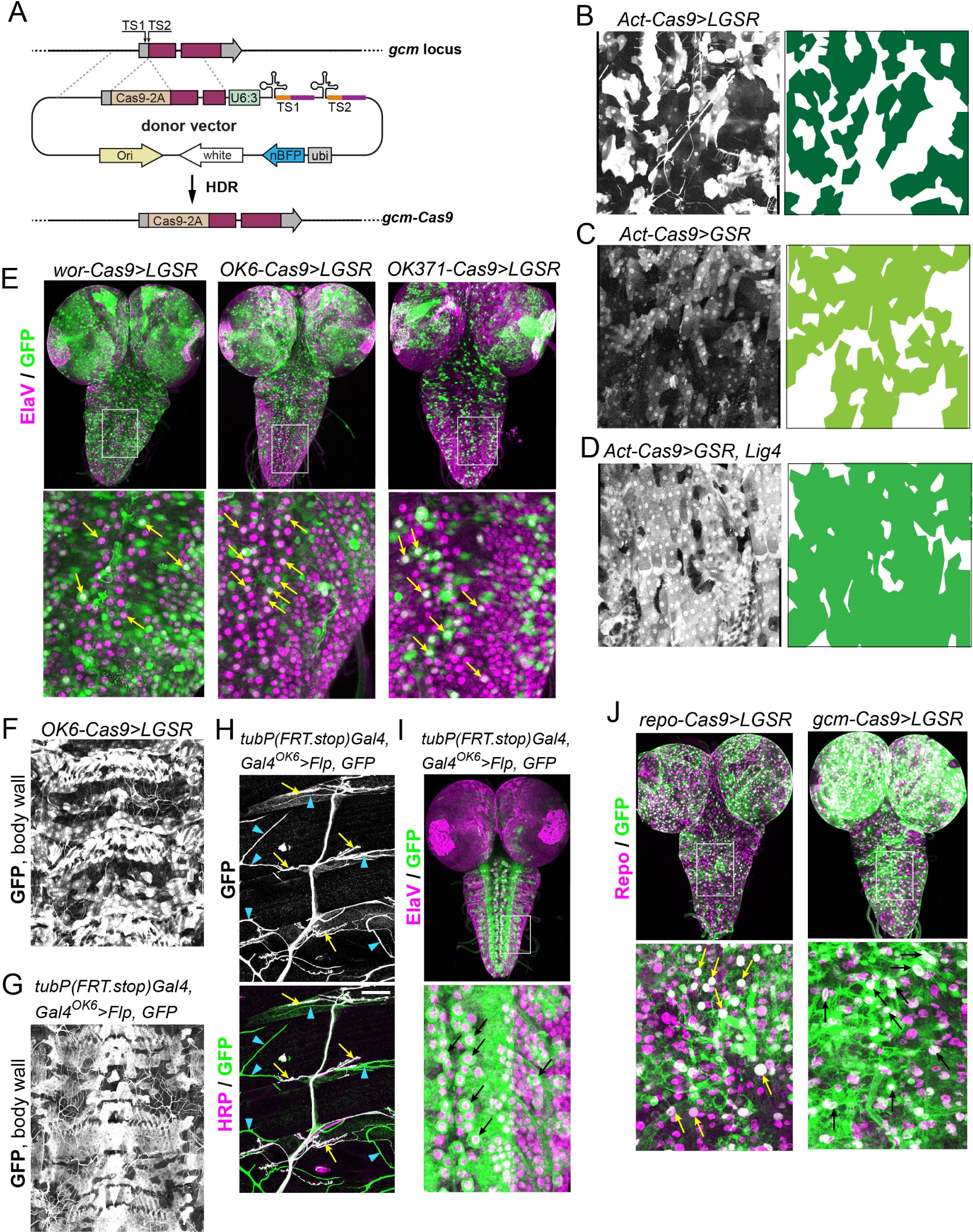
Cas9 activity patterns characterized by SSA reporters (related to Figure 1). **A**, a diagram of the generation of *gcm-Cas9* by CRISPR-mediated knock-in. A Cas9-2A coding sequence is inserted in-frame immediately after the start codon. **B-D**, activity pattens of Act-Cas9 in epidermal cells visualized by *LGSR* (B), *GSR* (C), and *GSR* in *lig4* mutant background (D). The panels on the right show masks of GFP-expressing cells. Images were taken using the same brightness setting. **E**, activity patterns of motoneuron-specific Cas9 in the larval brain visualized by *LGSR*. Elav antibody staining (magenta) shows neuronal nuclei. The lower panels are enlarged views of the boxed regions in the upper panels. Yellow arrows indicate *LGSR*-labeled neurons. **F**, activity pattern of *OK6-Cas9* in epidermal cells and trachea on the larval body wall, visualized by *LGSR*. **G-I**, activity patterns of *OK6-Gal4* in epidermal cells and trachea (G), motor neurons (H), and the larval brain (I), visualized by *tubP(FRT.stop)Gal4, UAS-Flp, GFP*. In (H), yellow arrows indicate GFP-labeled neurons and blue arrowheads indicate GFP-labeled trachea. **J**, activity patterns of glia-specific *repo-Cas9* (left panels) and *gcm-Cas9* (right panels) in the larval brain, visualized by *LGSR*. Larval brains were stained for the glial nuclear protein Repo (magenta). The lower panels are enlarged views of the boxed regions in the upper panels.

**Figure S2.**
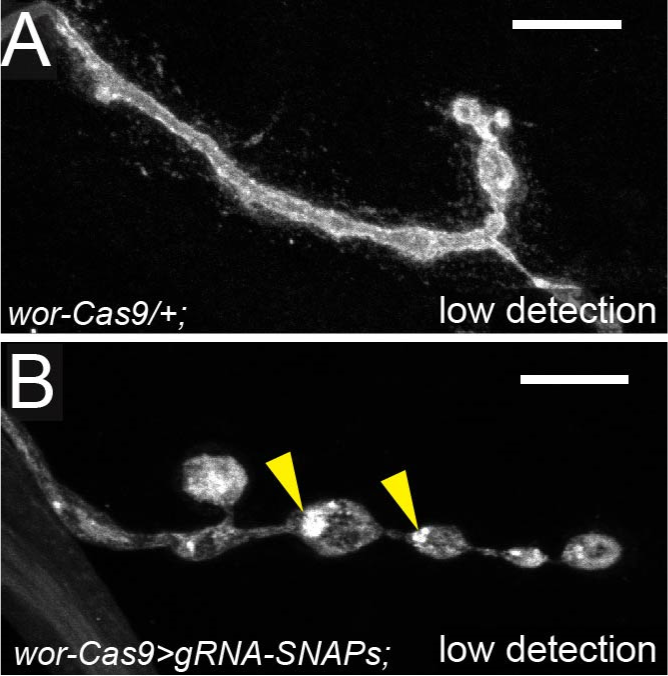
KO of SNAP genes causes intra-axonal membrane accumulation in boutons (related to Figure 3). **A** and **B,** NMJs of the control (A) and *Snap24/Snap25/Snap29* KO induced by *wor-Cas9* (B), imaged using a lower detection setting. Neurons are shown by HRP staining. Scale bar: 10μm. Related to Figure 3A-3B. Yellow arrowheads indicate dense puncta inside the presynaptic compartment.

**Figure S3.**
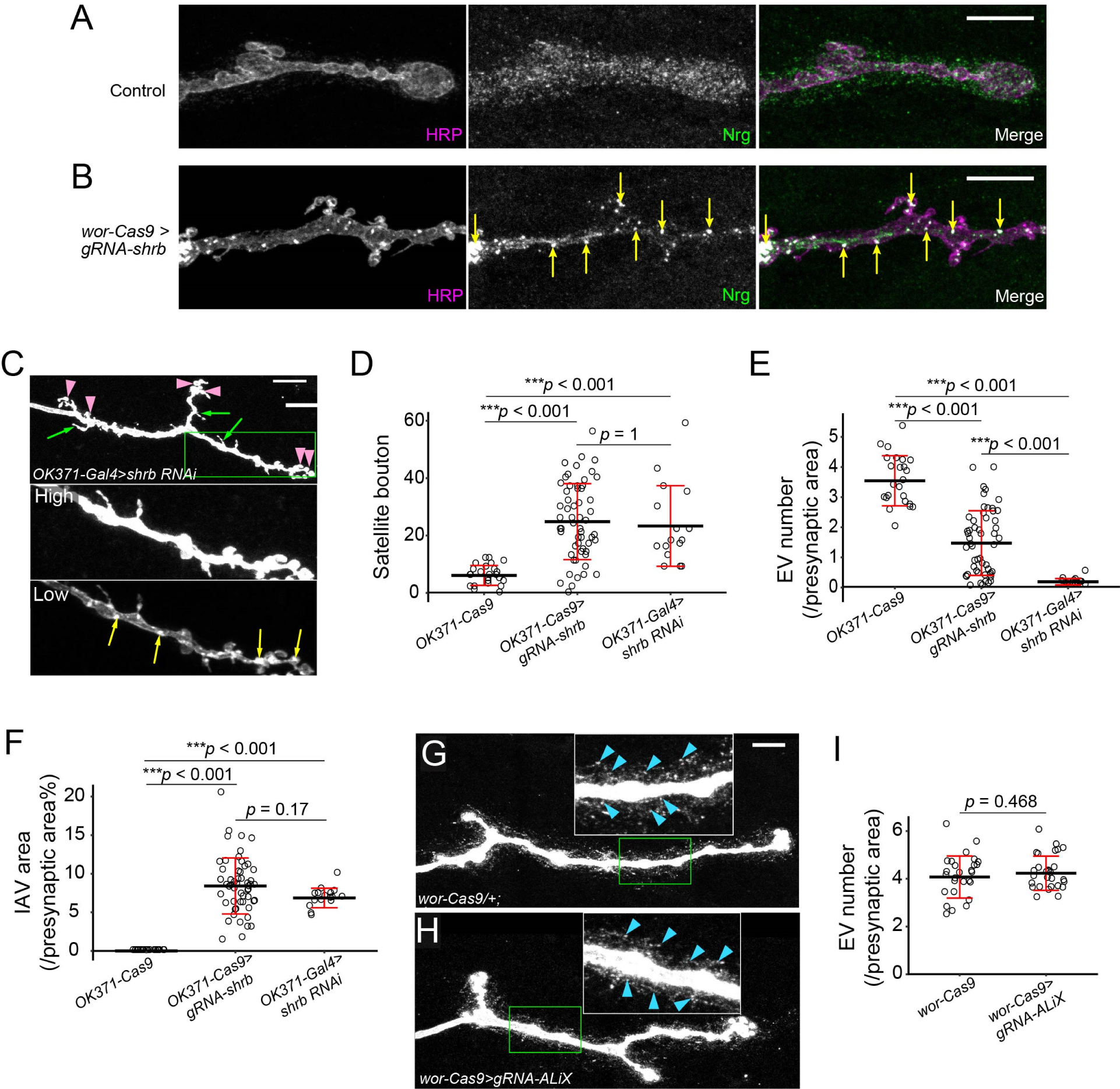
The KO phenotypes of ESCRT components at the NMJ (related to Figure 4) **A** and **B,** Nrg distribution at the NMJ of the control (A) and *shrb* KO by *wor-Cas9* (B). Axon membranes are visualized by HRP staining, and Nrg protein is detected by antibody stainining. Scale bar: 10μm. Yellow arrows indicate IAV colocalization with Nrg aggregation. **C**, neuronal-specific *shrb* KD induced by RNAi. “High” and “Low” panels show the zoomed-in view of the area enclosed by the green box imaged at high and low intensity settings. Pink arrowheads indicate satellite boutons, and green arrows indicate filamentous protrusions formed by presynaptic membrane. Yellow arrows indicate IAVs. Scale bar: 10μm. **D**, satellite bouton numbers in *shrb* KO and *shrb* KD motoneurons. One-way ANOVA, F(2,94) = 22.143, *p* = 1.38×10^−8^. Each circle represents an NMJ: *OK371-Cas9*, n= 24; *shrb^OK371-Cas9^*, n = 57; *shrb-RNAi^OK371-Gal4^*, n = 16; between-group *p* values are from multiple comparison test using Bonferroni adjustment. **E**, EV numbers normalized by the presynaptic area. One-way ANOVA, F(2,90) = 71.076, *p* < 2.2×10^−16^. Each circle represents an NMJ: *OK371-Cas9*, n= 24; *shrb^OK371-Cas9^*, n = 53; *shrb-RNAi^OK371-Gal4^*, n = 16; between-group *p* values are from multiple comparison test. **F**, IAV areas normalized by the presynaptic area. One-way ANOVA, F(2,91) = 75.239, *p* < 2.2×10^−16^. Each circle represents an NMJ: *OK371-Cas9*, n= 24; *shrb^OK371-Cas9^*, n = 54; *shrb-RNAi^OK371-Gal4^*, n = 16; between-group *p* values are from multiple comparison test. **G** and **H,** NMJ morphology in the control (G) and *ALiX* KO (H) motoneurons. Neuronal membrane and EVs are visualized by HRP staining. Inset: zoomed-in view of the area enclosed by the green box. Blue arrowheads indicate the EVs surrounding the presynaptic compartment. **I**, EV numbers normalized by the presynaptic area. t-test, t(50.1) = −0.73, *p* = 0.468.

**Figure S4.**
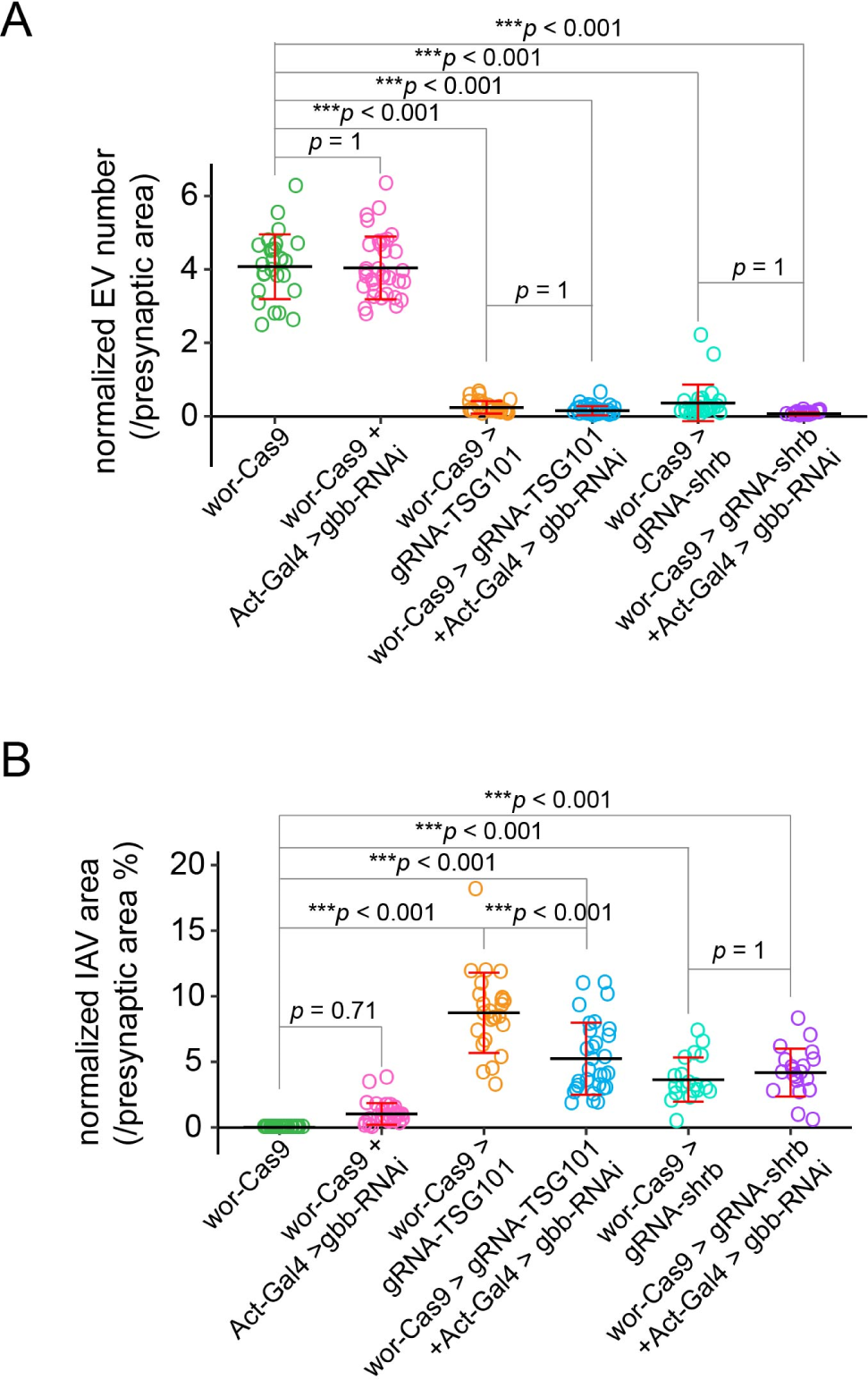
The impacts of *gbb* KD on EV numbers and IAV areas (related to Figure 5) **A**, normalized EV numbers in genotypes represented by Figure 5***A***–***F***. One-way ANOVA, F(5,158) = 349.56, *p* < 2.2×10^−16^. Each circle represents an NMJ: *wor-Cas9*, n= 27; *gbb-RNAi^Act-Gal4^*, n = 34; *TSG101^wor-Cas9^*, n = 26; *TSG101^wor-Cas9^ / gbb-RNAi^Act-Gal4^*, n = 30; *shrb^wor-Cas9^*, n = 25; *shrb^wor-Cas9^ / gbb-RNAi^Act-Gal4^*, n = 22; between-group *p* values are from multiple comparison test using Bonferroni adjustment. **B**, normalized IAV area in genotypes represented by Figure 5***A***–***F***. One-way ANOVA, F(5,148) = 67.193, *p* < 2.2×10^−16^. Each circle represents an NMJ: *wor-Cas9*, n= 27; *gbb-RNAi^Act-Gal4^*, n = 33; *TSG101^wor-Cas9^*, n = 25; *TSG101^wor-Cas9^ / gbb-RNAi^Act-Gal4^*, n = 30; *shrb^wor-Cas9^*, n = 19; *shrb^wor-Cas9^ / gbb-RNAi^Act-Gal4^*, n = 20; between-group *p* values are from multiple comparison test using Bonferroni adjustment. The datasets of *wor-Cas9>gRNA-shrb* and *wor-Cas9>gRNA-TSG101* are the same as in Figure 4.

